# Simulating The Effect of the Radiotherapy on Cancer Stem Cell Evolution

**DOI:** 10.1101/2025.01.23.634433

**Authors:** Yusri Dwi Heryanto, Seiya Imoto

## Abstract

Radiotherapy is one of the most effective treatments for cancer. However, the radiation on cancer often leads to further tumor evolution and resistance. Accumulating evidence suggests that the cancer stem cells (CSCs) play a significant role in the tumor’s radioresistance and invasiveness. Therefore, we set out to investigate the aggressiveness and radioresistance that emerge because of radiation-induced selective pressures on CSCs. We developed an agent-based model (ABM) that tracks the evolution and growth of the cancer after treated by different regimens of radiotherapy. In the ABM, each cell is an agent that follows some predefined rules for their behaviors and interactions with other agents. We showed that the radiotherapy treatment increased the resistance, proliferation rate, and the CSC proportion of the tumor. Our simulation also showed that the CSC-targeted treatment is important for eliminating tumor completely.

## Introduction

Radiotherapy is one of the most common cancer treatments. About half of the cancer patients will get radiotherapy at some point during their therapy^1^. Ionizing radiation (IR) kills cancer cells by the direct disruption of DNA or by the reactive oxygen species indirect DNA damage^2,3^. The treatment is usually given in fractionated irradiation to limit the damage to normal tissues. The most common types of fractionation are: conventional fractionation (2Gy per day for 30 days), hyperfractionation (fractional doses smaller than 2Gy, given 2 or 3 times daily), and hypofractionation (fractional doses more than 2Gy with fewer fractions than conventional).

However, the cancer cells can adapt and develop resistance during IR therapy^4^. In recent years, there is accumulating evidence on the existence of a rare tumor subpopulation known as cancer stem cells (CSCs) that plays an important role in radioresistance^5,6^. CSCs are defined as cancer cells that possess the capacity to self-renewal, develop to heterogeneous lineages of cancer cells, and proliferate indefinitely^7^. CSCs are important in the treatment planning because it less sensitive to the therapy^6,8^and promote cancer invasiveness^9,10^. Paradoxically, the treatment itself can induce mutagenesis and selective pressures that will increase tumor resistance and invasiveness^11–13^. Failing to eliminate CSCs during treatment can lead to tumor relapse^14^.

The study of *in vivo* behavior of CSCs is important to understand the mechanism of tumor aggressiveness and resistance. However, there are many challenges for CSCs *in vivo* observation, such as the limitations of current live imaging techniques and the reliability of the CSCs biomarker^15,16^. The computational models can help to simulate *in vivo* the complex behavior and interaction of CSCs where they are difficult to observe in the wet lab experimental research.

We present an *in silico* agent-based model (ABM) to investigate the evolutionary trajectory of the CSCs that contribute to the tumor aggressiveness and resistance after radiotherapy. Agent-based model is a computational model for simulating the reaction of autonomous agents to their environment or other agents according to a predefined set of rules. We can describe a simple ABM as follows: agents may receive input from their environment or other agents, agents provide output to their environment or other agents, and then the agents react based on the input they received. The ABM allows researchers to study how system level properties emerge from the individual’s actions, as well as how the system affects individuals. ABM has been used in many CSCs studies to investigate the effect of migration^17^, plasticity^18^, mutation^19^, and therapy^20^.

In our experiment, we simulated and followed the temporal evolution of the cancer cells traits, such as the capability of proliferation, migration, symmetric division, plasticity, and radioresistance after IR treatments. We showed that the IR treatment lead to the increase of proliferation capability, resistance, symmetric division, and CSCs proportion. These will promote the tumor aggressiveness, resistance, and relapse post-treatment. Our simulation also suggested that the CSCs targeting-treatment is essential to the tumor complete elimination.

## Results

We used an ABM to simulate the growth and evolutionary dynamics of the tumor under various radiotherapy regimens. In ABM, each tumor cell, either cancer stem cell (CSC) or non-stem cancer cell (CC), was represented by an agent. On every time step, each agent was selected in random order. The selected agent then reacted following predefined rules shown in Fig.1. In brief, the agent attempted the replication following the rules in the replication pathway. If the agent had a deleterious mutation, the agent became unviable. The deleterious mutation was characterized by a negative value in one or more cell traits (e.g. proliferation rate = − 0.01). Then, the agent checked its neighboring space. If there was no empty space, the agent was entering the quiescent state. If there was an empty space, the agent was going to replicate with probability *p*_*prol*_. On replication, the agents may spontaneously die, mutate, or change its phenotype (i.e. CSC become CC or CC become CSC). In our experiment, we simulated the mutation on the traits proliferation rate, migration rate, symmetric division probability, phenotype change probability, and the radioresistance. If the agent did not replicate, it might move to an empty space with probability *p*_*mig*_. After following the rules in the replication pathway, the agents followed the rules in the treatment pathway. If the agent were irradiated at the current step, it may die depends on its resistance and the radiation dosage. The unviable cells and death cells were removed from the simulation.

**Figure 1.**
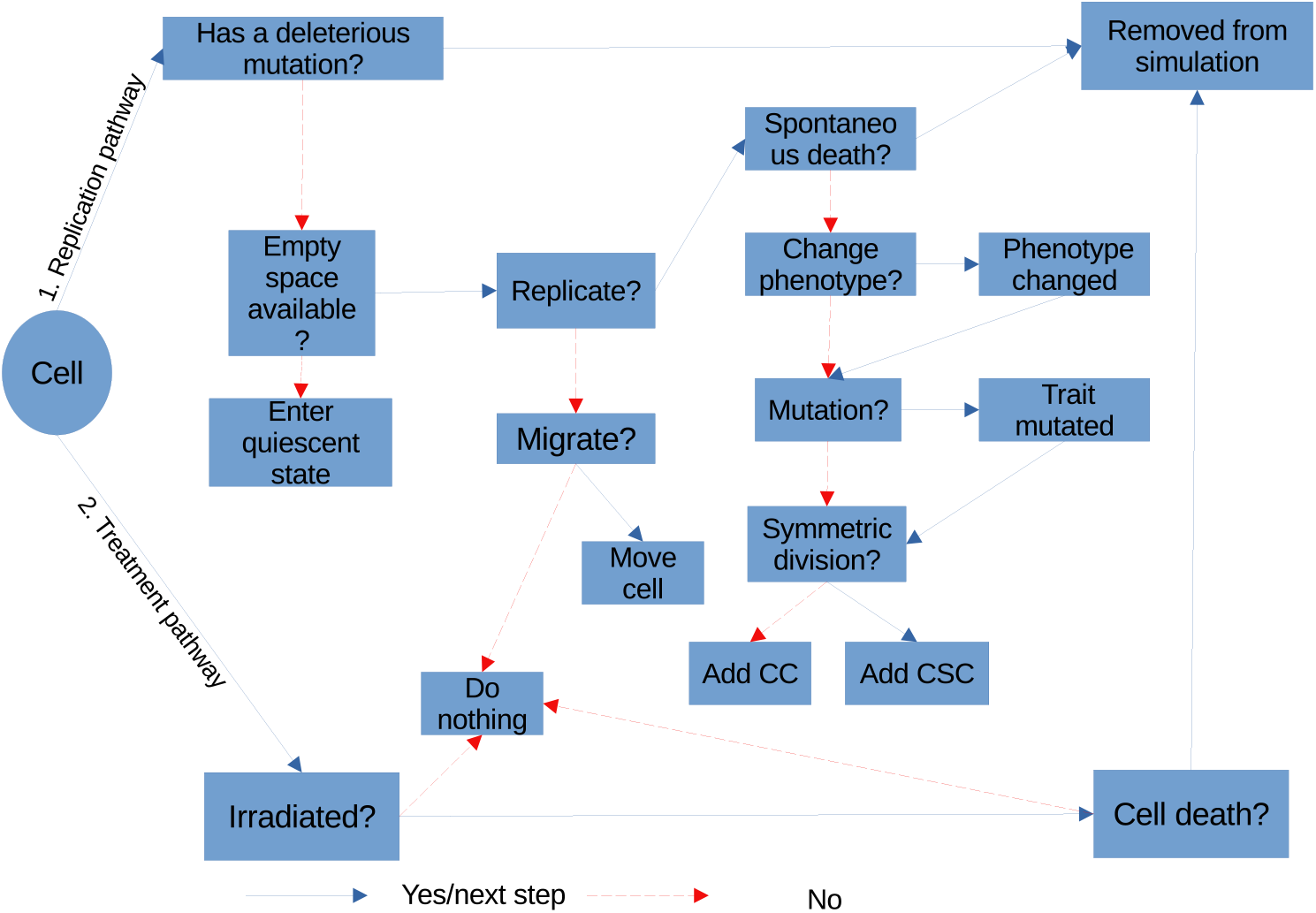
The flowchart of the ABM rules. First, the agents follow the rules on the replication pathway that control the behavior of the agents when they are attempting to replicate. Next, they follow the rules in the treatment pathway, where the agents may get irradiated and die.

We started the simulation at the time step *t* = 0 and started the treatment on time step *t* = 1 with one of 4 regimens: control (ie. no treatment), conventional radiotherapy, hyperfractionated radiotherapy, and hypofractionated radiotherapy. Each time step was equal to 1 hour. We also simulated CSC-targeted radiotherapy that used radiosensitizer to lower the CSCs resistance. The treatment finished at *t* = 720 or after 30 days. From the *t* = 0 until *t* = 1200, the evolution of proliferation, symmetry division, migration, dedifferentiation, and resistance traits were recorded. We also recorded the cancer cell number, CSCs ratio, and quiescent cell ratio at each time step.

We did not find any significant effect of regimens to the evolutionary dynamics of the tumor traits. However, the mutation rate affected the cell traits during therapy.

### Increased resistance during therapy

When irradiated, the cell may die with the probability 1 −*SF*(*D, ξ, λ*) where *SF* is a linear quadratic surviving fraction formula^21^ that has inputs radiation dose *D*, CSC resistance factor *λ*, and quiescent resistance factor *ξ*. The lower resistance factor *ξ* and *λ* means the lower the probability the cell to die during IR treatment(i.e. increased resistance). We set the CSCs to have lower *λ* and the quiescent cells have lower *ξ* because they are inherently more radioresistant than other cancer cells.

On the middle graph of Fig. 2, we observed that the mean resistance factor *λ* of the tumor cells decreased during the radiotherapy (i.e. increased radioresistance). However, the resistance approached the control value after the treatment finished at *t* = 720. We also observed that the CSC ratio graph negatively correlated with the resistance factor *λ*. This result suggests that the mean resistance increases mainly because of the increase of CSC ratio in the tumor. CSCs which were inherently radioresistant were more likely to survive and contributed to the radioresistance of the tumor than the CCs. The acquired radioresistance during repeated irradiation were confirmed in experimental studies^4,12^. This result is also consistent with an *in vitro* study by Fukui *et al*. that investigated the characteristic of cells after radiation^22^. They found that the tumor that irradiated to daily 2Gy of X-rays and became radioresistant had higher fractions of aldehyde dehydrogenase (ALDH) positive CSCs than the control tumor. They suggested that the acquired radioresistance after irradiation was intrinsically related to the ALDH-positive CSC subpopulation of tumor cells.

**Figure 2.**
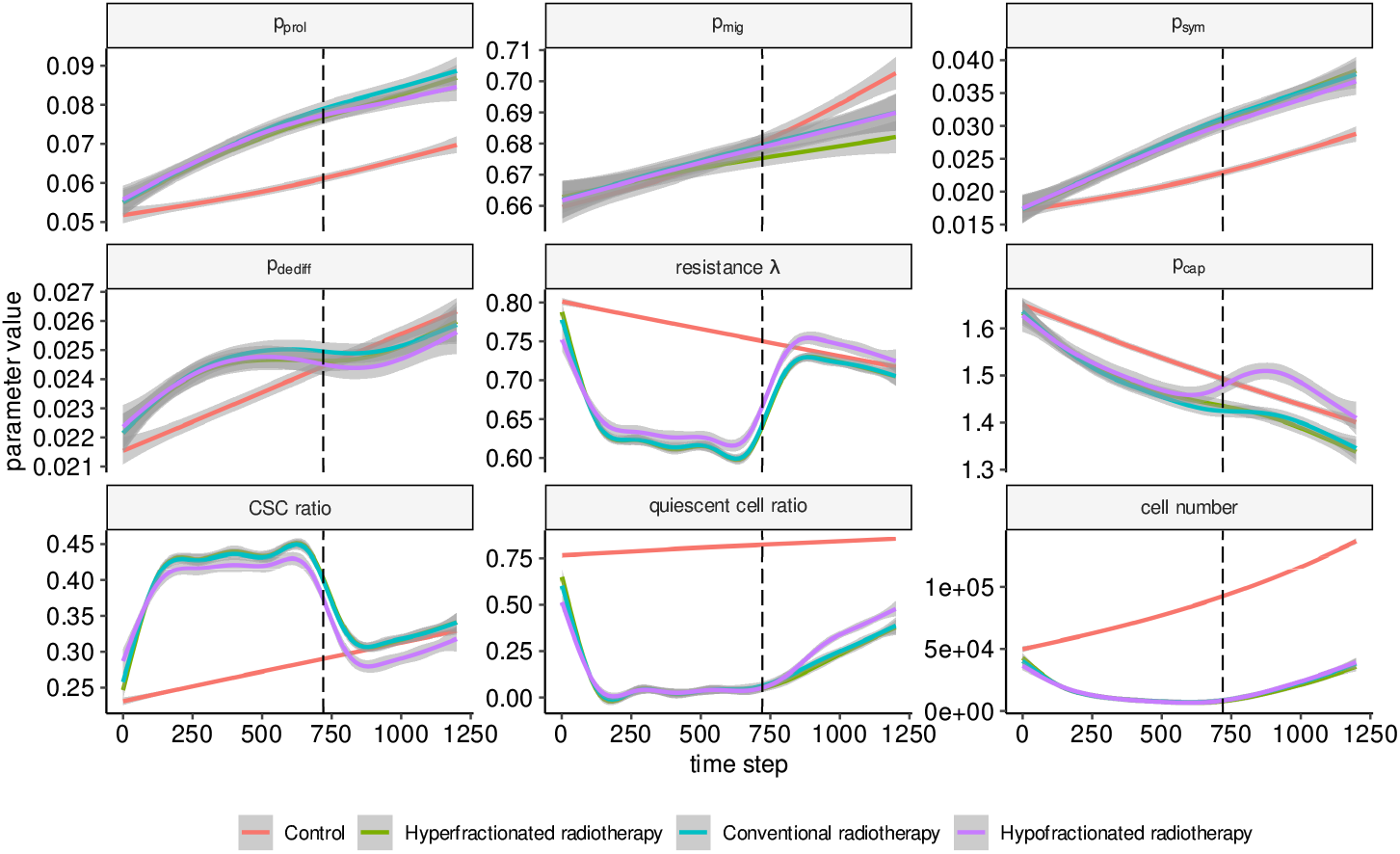
The evolution of the mean of traits proliferation rate *p*_*prol*_, migration rate *p*_*mig*_, symmetric division probability*p*_*sym*_, phenotype change probability*p*_*dedi f f*_, and resistance factor *λ* from the tumor during and after various radiotherapy regimens. We recorded the traits from *t* = 0 until *t* = 1200. The therapy started at *t* = 1 and finished at *t* = 720 (black dashed line). We used *p*_*mut*_ = 0.5 on this experiment. The 0.95 level of confidence interval (shaded area) was calculated from 20 replicates of experiments. We also recorded the proliferation capacity *p*_*cap*_ of the tumor, CSC ratio, quiescent cell ratio, and cell number.

The resistance of the treated tumors was back to the control value after the treatment finished. It might be caused by the normalization of the CSC ratio after treatment.

### Increased proliferation rate during therapy

We found that the proliferation rate was increasing *p*_*prol*_ faster than the control during the treatment of every radiotherapy regimen (upper left Fig. 2). In our model, the cancer cells that have a high proliferation rate were more likely to survive and passed their traits to their descendant. There is strong evidence for accelerated repopulation during fractionated radiotherapy from many experiments^23–26^. The mechanisms that underlie accelerated repopulation during radiotherapy are not well understood. The improvement of nutritional and oxygen status and the activation of EGFR have been proposed as the cause of the accelerated repopulation^27,28^. We showed that the accelerated repopulation might be due to the high proliferation rate cells that survived radiation-induced selective pressure.

### CSCs proportion change

During the treatment, the ratio of CSC to the total number of the cells was increasing (lower left of Fig. 2). These result is consistent with the clinical and experimental findings^22,29,30^. The elimination of non-radioresistant CCs is the main cause of the increasing ratio of the CSCs. We found that the symmetric division probability also increasing. The CSCs divide symmetrically to make two CSC daughter cells, while asymmetric cell division produces one CSC and one CC daughter cells. The phenotypic change probability was also increased during the treatment. Put together, the radiotherapy created a condition that increases the CSCs ratio in the tumor by eliminating the non-radioresistant CCs and by promoting the survival of cell that have high phenotypic change or symmetric division probability.

### Effect of mutation rate on tumor growth during therapy

We simulated the effect of mutation rate on growth and evolutionary dynamics of the tumor (Fig. 3, Fig. S1). Each trait mutates with the probability *p*_*mut*_ on replication. We observed the effect of *p*_*mut*_ = (0.1, 0.3, 0.5) on 2 groups of tumor: control tumor (i.e. without treatment) and tumor treated with conventional radiotherapy (i.e. radiation dose 2Gy daily for 30 days). We found that the higher mutation rate tumors have a bigger number of cell during and after therapy, as shown in the cell number graph on Fig. 3. The similar finding has been reported by Poleszczuk *et al*.^19^. On the traits *p*_*prol*_, *p*_*sym*_, and *p*_*dedi f f*_, the tumors with high mutation rate have a bigger deviation from the control. From these findings, we hypothesized that the higher mutation rate is beneficial to the tumor to survive and adapt in the hostile condition. Radiation on the tumor with high mutation rate will be more likely to select the cells that have higher proliferation rate, symmetric division probability, and phenotypic change probability to survive.

**Figure 3.**
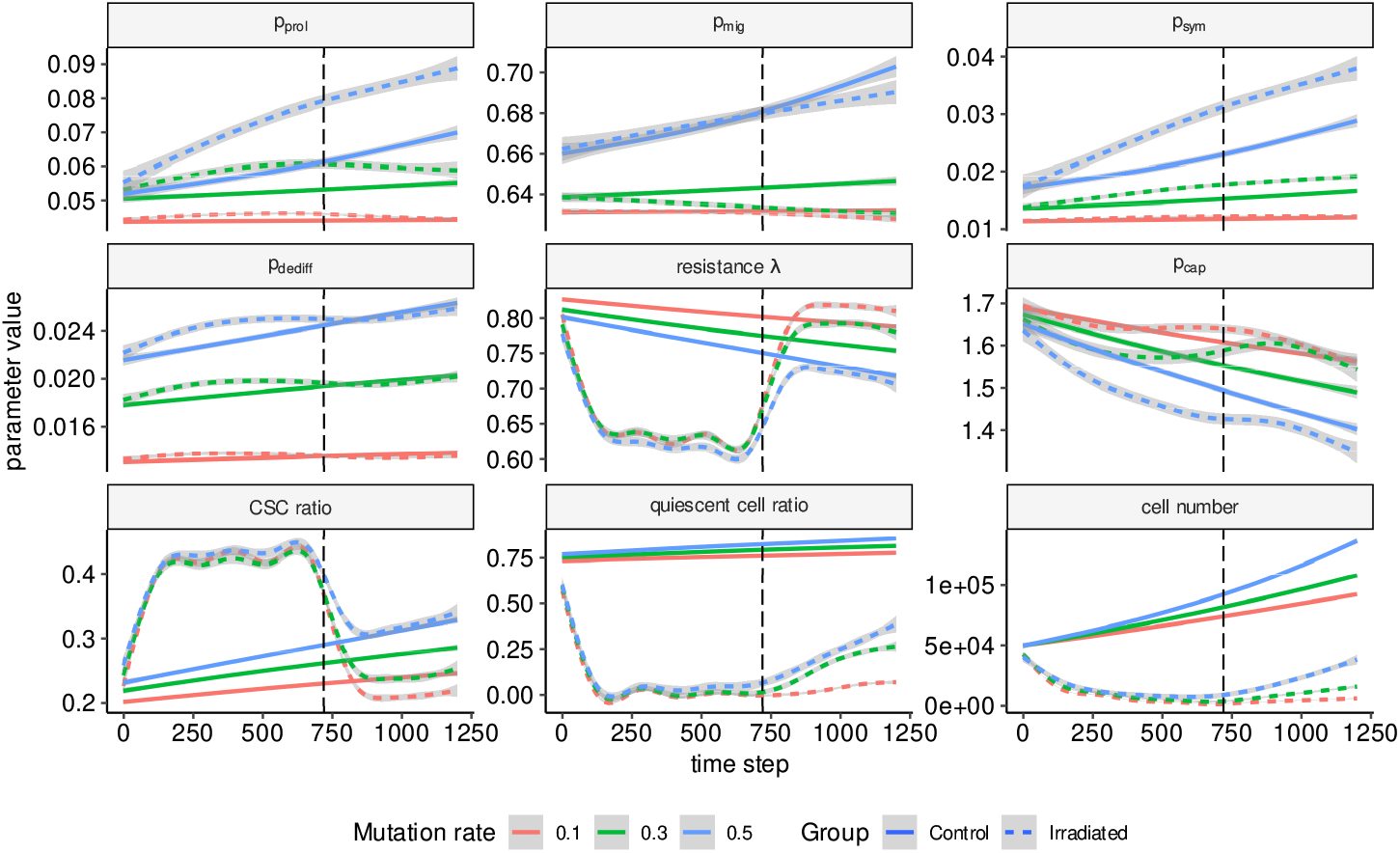
The evolutionary and growth dynamics of the tumor treated with conventional radiotherapy (2Gy per 24 hours for 30 days) under different mutation rates.

However, the resistance of the treated tumors on each mutation rate was not too much different during treatment. As stated before, it suggests the resistance change was mainly due to the change of CSCs ratio rather than the effect of the mutation.

### CSC-targeting treatment is essential to tumor elimination in the simulation

Our findings in this experiment suggest that CSCs are the major culprits of tumor development and therapeutic resistance. Thus, a treatment that target CSCs might increase the efficacy of currently available treatment regimens and reduce the chance relapse. We simulated a CSC-targeted-therapy that combines a CSC-targeting-radiosensitizer and conventional radiotherapy. Radiosensitizers are compounds that increase the effectiveness of radiotherapy in killing tumor cells^31^. We assumed that the CSC-targeting-radiosensitizer increased the resistance factor *lambda* of the CSCs by 0.5. From the 20 tumors treated with targeted radiotherapy, only one tumor survived until *t* = 1200. Most of the tumors were eliminated before *t* = 600 (i.e. days 25). In contrast, all tumors in control and conventional radiotherapy group were survived (Fig. 4, Fig. S1).

**Figure 4.**
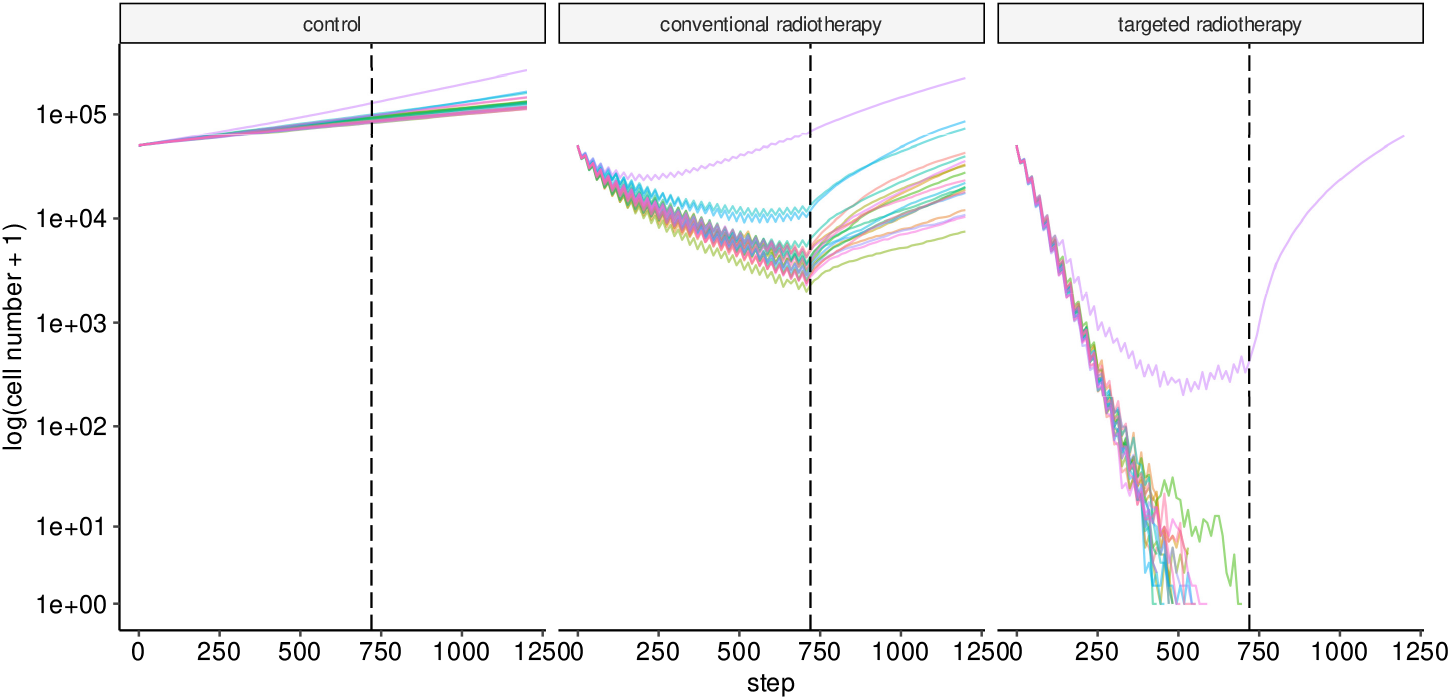
The comparison of tumor growth on control, conventional radiotherapy, and targeted therapy group. The black dashed line on *t* = 720 marks the day when the treatment was finished. The *y-axis* is the cell number under *log*(1 + *x*) transformation.

**Figure 5.**
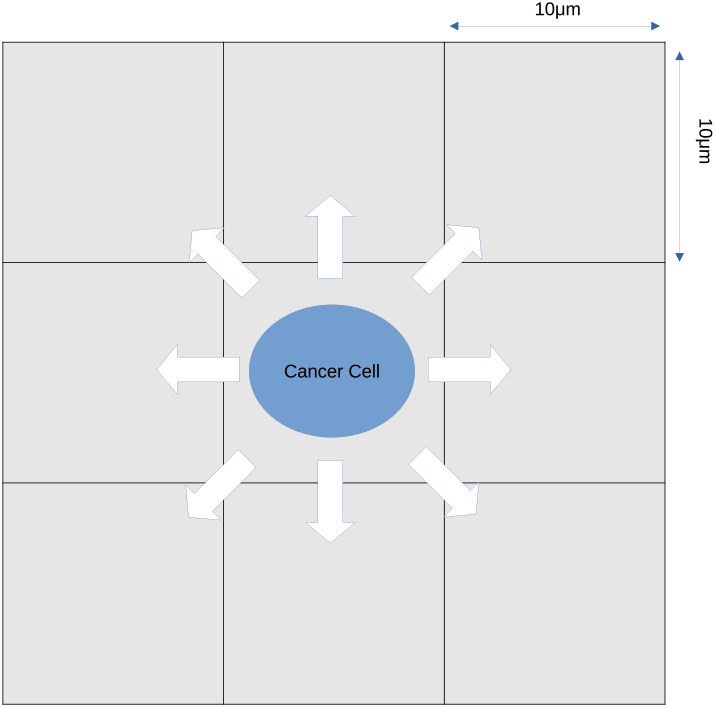
An agent occupies a 10 µm × 10 µm grid in lattice grid space. The agent can check its 8 neighboring grid for an empty space.

## Discussion

To study the *in vivo* behavior of CSCs, we need a specific CSC biomarker. However, we have not yet found the CSC ‘best marker’ that is entirely reliable and truly CSC-specific^16^. The limitation of resolution and depth of current imaging technology also become challenges^15^. It is difficult, therefore, to observe and draw general conclusions about *in vivo* CSC behaviors.

To overcome these limitations, we used ABM to observe in real time the evolutionary dynamics of the cancer cells, especially CSCs. ABM is suitable for CSCs research because it enables us to monitor, albeit *in silico*, the tumor growth, cellular distribution, and genetic mutations in real time. By doing so, we can observe the emerging behavior of the CSCs from the interaction of many traits and conditions. The ABM can be reproduced and extended easily. The results of the ABM have strong mathematical support because we can see the ABM as an axiomatic system that produces theorems. Of course, the empirical validity depends on the assumptions and the validity of the model. As other simulation study, the model also needs to be verified in the experimental study.

We found some interesting findings in this study. First finding is the CSC ratio affects the radioresistance more than the mutation that causes *de novo* radioresistance. The resistance of the low mutation rate and high mutation rate tumor were not so different during the treatment. However, the resistances were negatively correlates with the decreasing resistance factor *λ* (i.e. increasing radioresistance). The increasing radioresistance during and after the radiotherapy are well documented^4,12^ and multiple mechanisms to explain it have been proposed^32^. We showed the CSC might be the primary cause of the radioresistance. This finding has implication in cancer therapy if it is validated. Rather than targeting the radioresistance-related genes to overcome resistance, it might be better to target the CSCs that cause radioresistance. The heterogeneity of gene expression in a tumor leading to multiple, concurrent resistance mechanisms made it difficult to target a single gene^33^. Moreover, the radioresistance genes on non-stem cancer cells were more likely to be lost from the population due to limited proliferation capacity^34^. Thus, targeting the radioresistance genes might be suboptimal to control radioresistance. However, we should note that our model did not consider the tumor microenvironment, vascularity, hypoxia, and size, which are also affecting the resistance^32,35^. Our model started with the tumor with about 50000 cells and the diameter approximately 2000µm. It has been demonstrated that the tumor with the diameter about 2000µm or less is in the avascular phase where cancer cells can get the nutrient merely by diffusion^36^. Thus, we can ignore the effect of vascularity, hypoxia, and size in our model.

We also found that the cell with the high proliferation rate, high symmetric division probability, and high phenotypic change probability were more likely to survive and passed the traits to their descendant. Increased proliferation correlates strongly with poor prognosis and higher probability of relapse^37–39^. High symmetric division and phenotypic change probability contribute to increasing the CSC ratio. Altogether, these traits from the cells that survive made the tumor more aggressive and resistant.

Because of its critical role in tumor invasiveness, resistance, and treatment failure, many researchers suggested that the targeting of CSC is an effective way to eradicate cancer^40,41^. Our finding supported the idea of CSCs targeting for cancer elimination. From all treatment simulations, only the CSC targeting regimen can totally eradicate the tumor. Here, we used a radiosensitizer that can specifically target the CSCs. In reality, there are multiple problems that need to be solved to eliminate CSCs. First, the characteristics of CSCs in many tumors are not well defined^42^. Second, currently there is no reliable and specific biomarker and drug that can target CSCs^16^. Third, a tumor can have a diverse population of CSCs^43^.

The model presented here is a complex but elementary approach to provide an evolutionary explanation of the emerging tumor resistance and aggressiveness during radiotherapy. Our current model still has many limitations. We limited our study to 2-dimensional space due to memory and time limitation. A previous study found that the 2-dimensional ABM qualitatively mimic 3-dimensional ABM CSCs dynamics^44^. Our model, like other simulation models, is subject to gross oversimplification. We did not include the microenvironment factor, immune status, and vascularity. However, it might help the researchers to observe phenomenon which are difficult to observe in real experiment. It may provide novel and interesting insight. This model can be further developed by including environment factor, CSC-targeting strategy, immune cell interaction, and other treatment modality. Integrating mathematical and experimental models is needed to understand the complex behavior of CSCs.

## Methods

We developed an ABM to investigate the evolution of cancer cells phenotypic after IR treatments. An agent, either CSCs or non-stem cancer cells (CC), occupies a 10 µm × 10 µm grid in a 1000 × 1000 2-dimensional lattice grid 5. Each cancer cell is defined as an agent with properties listed in Table 1.

**Table 1.**
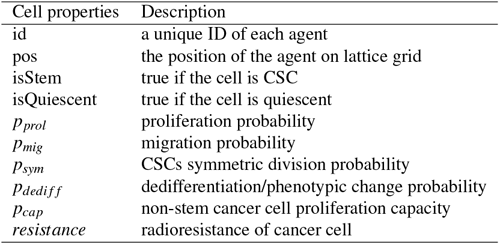
The properties of each agent.

**Table 2.** The value of parameters when initiating the model.

| Parameters | Starting value | Notes |
| --- | --- | --- |
| isStem | true |  |
| isQuiescent | false |  |
| $p_{prol}$ | 1/24 | 1 proliferation per day |
| $p_{mig}$ | 15/24 | 150 $\mu\text{m}$ per day |
| $p_{sym}$ | 0.01 | |
| $p_{dediff}$ | 0.01 | |
| $p_{capacity}$ | 15 | |
| $\lambda_{CSC}$ | 0.1376 | the resistance factor of CSC |
| $\lambda_{CC}$ | 1.0 | the resistance factor of CC |
| $\xi_{quiescent}$ | 0.5 | the resistance factor of quiescent cell |
| $\xi_{non-quiescent}$ | 1.0 | the resistance factor of non-quiescent cell |
| $p_{death}$ | 0.01 | |
| $\sigma_{prol}$ | 0.01 | SD proliferation parameter |
| $\sigma_{mig}$ | 0.05 | SD migration parameter |
| $\sigma_{sym}$ | 0.005 | SD symmetry parameter |
| $\sigma_{dediff}$ | 0.005 | SD dedifferentiation parameter |
| $\sigma_{resistance}$ | 0.05 | SD resistance parameter |

The simulation time step is 1 hour. Thus, 24 time steps equal to one day. At each step, we selected each of the agent on random order and updated their behavior. First, they need to do steps in replication pathway. In replication pathway, the agents may replicate, change phenotype, migrate, mutate, enter quiescent state, or die spontaneously. After replication pathway, we started the IR treatment simulation in treatment pathway. In treatment pathway, the cell may die and removed from simulation due to irradiation damage (Fig. 1).

A non-stem cancer cell (CC) has a limited proliferation capacity, *p*_*cap*_, due to telomere shortening^45^. Every time the CC mitose in our model, the proliferation capacity is reduced by one (i.e. *p*_*cap*_ −1) in each daughter to reflect the telomere shortening. In contrast, the CSCs have unlimited proliferation capacity^7^. Thus, the *p*_*cap*_ of the CSC parent will be passed to the daughters without any reduction. If the *p*_*cap*_ = 0, the cell will undergo cell death in the next replication attempt. A cell need at least one adjacent empty space for replication. If there is no neighboring empty space (i.e. none of the adjacent 8 grids is vacant), the cell go into quiescent state (i.e. *isQuiescent* parameter become true) until a vacant space available again^46^. In each division, a CSC has probability *p*_*sym*_ to perform symmetric division where the both daughters is CSC and 1 −*p*_*sym*_ to perform asymmetric division where the daughters are one CSC and one CC^47^. An CC can dedifferentiated to CSC with the probability *p*_*dedi f f*_ on each division attempt^48^. *Vice versa*, an CSC may loss their stem-like properties and become CC with the same probability, *p*_*dedi f f*_. When the cell changed their phenotype, their resistance also changed based on their new phenotype. If the cell do not replicate, it has the *p*_*mig*_ chance to migrate to an adjacent empty space. At each proliferation attempt step, we assumed the spontaneous apoptosis has the probability *p*_*death*_ to occur randomly^49^ on any cell. To investigate phenotypic evolution, we model mutation events in each of cancer cell. Each of the parameters *p*_*prol*_, *p*_*mig*_, *p*_*sym*_, *p*_*dedi f f*_, and *resistance* are subject to mutation events with the probability *p*_*mut*_. We set the mutation rate *p*_*mut*_ = 0.5 to observe the effect of the different regimens of radiotherapy to the traits evolution. To investigate the effect of the mutation rate, we treated 3 group of tumors: low, medium, and high mutation rate with *p*_*mut*_ are equal to 0.1, 0.3, 0.5, respectively. Then, we treated these 3 groups with conventional radiotherapy or no treatment/control. We assume that a mutation event induces a stochastic change that follow a normal distribution where the mean of the distribution is the parent current parameter value. The *σ*_*prol*_, *σ*_*mig*_,*σ*_*sym*_,*σ*_*dedi f f*_, *σ*_*resistance*_ are the standard deviation (SD) for proliferation, migration, symmetry, dedifferentiation, and resistance parameters. If a parameter become negative, we assume the cell is unviable and removed from simulation.

The simulation was initialized with a CSC with the proliferation rate *p*_*prol*_ = 1*/*24, migration rate *p*_*mig*_ = 15*/*24, probability of symmetric division *p*_*sym*_ = 0.01, probability of phenotype changes *p*_*dedi f f*_ = 0.01, proliferation capacity *p*_*cap*_ = 15, and resistance *λ* = 0.1376. We grew the tumor until the cell number reached at least 50000. The mutation events will be triggered with the probability *p*_*mut*_ = (0.1, 0.3, 0.5) for traits *p*_*prol*_,*p*_*mig*_,*p*_*sym*_,*p*_*dedi f f*_, and *λ* on each replication attempt. We selected the parameters value based on previous studies^18,19^.

After we grew the tumor, we set *t* = 0 and treated the tumor from *t* = 1 until *t* = 720. We simulated 4 types of radiotherapy regiments: the conventional radiotherapy with 2Gy radiation per fraction applied every 24 hour, the hyperfractionated radiotherapy with 1Gy radiation per fraction applied every 12 hour, the hypofractionated radiotherapy with 4Gy radiation per fraction applied every 48 hour, and the targeted radiotherapy with the dosage same as conventional but we specifically targeted the CSCs nad lower its resistance. We calculated the surviving fraction treated with *D* dose of radiation using the linear quadratic formula *SF*(*D, ξ, λ*):

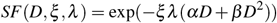

The parameters *ξ* and *λ* describe the radioresistance of quiescent cell and CSC respectively. The lower the *ξ* and *λ* is, the higher the resistance of the cell. The *ξ* of quiescent cells is generally lower than the non-quiescent cells^50^. The values of *ξ* is 0.5 and 1.0 for the quiescent and non-quiescent cell respectively. The *λ* of CSCs is lower than their CCs counterpart^6^. The *λ* value are influenced by mutation. The initial values of the *λ* are 0.1376 and 1.0 for CSCs and CCs, respectively. In the targeted radiotherapy, we increased the CSCs *λ* by 0.5. We fixed the parameters *α* = 0.3859 and *β* = 0.01148. The values of *ξ, λ, α*, and *β* were in line with previous studies^21^.

We followed and recorded the traits evolution from *t* = 0 until *t* = 1200. The cell numbers, the CSC to total cell number ratio, and the quiescent cell to total cell number ratio were also recorded. For each regiments, we replicated our experiment 20 times.

## Author contributions statement

Y.D.H. was responsible for the study conceptualization, the simulation, analyses, and visualization, and writing the original draft of the manuscript. S.I. was responsible for the funding acquisition, project administration, supervision, and editing the manuscript. All authors reviewed the manuscript.

## Additional information

## Data availability

The data in this study were generated using the source code at the GitHub repository (https://github.com/yusri-dh/CSCABM).

## Code availability

The source code to perform and replicate the simulation in our study is available at the GitHub repository (https://github.com/yusri-dh/CSCABM).

## Competing interests

The authors declare no competing interests

**Figure S1.**
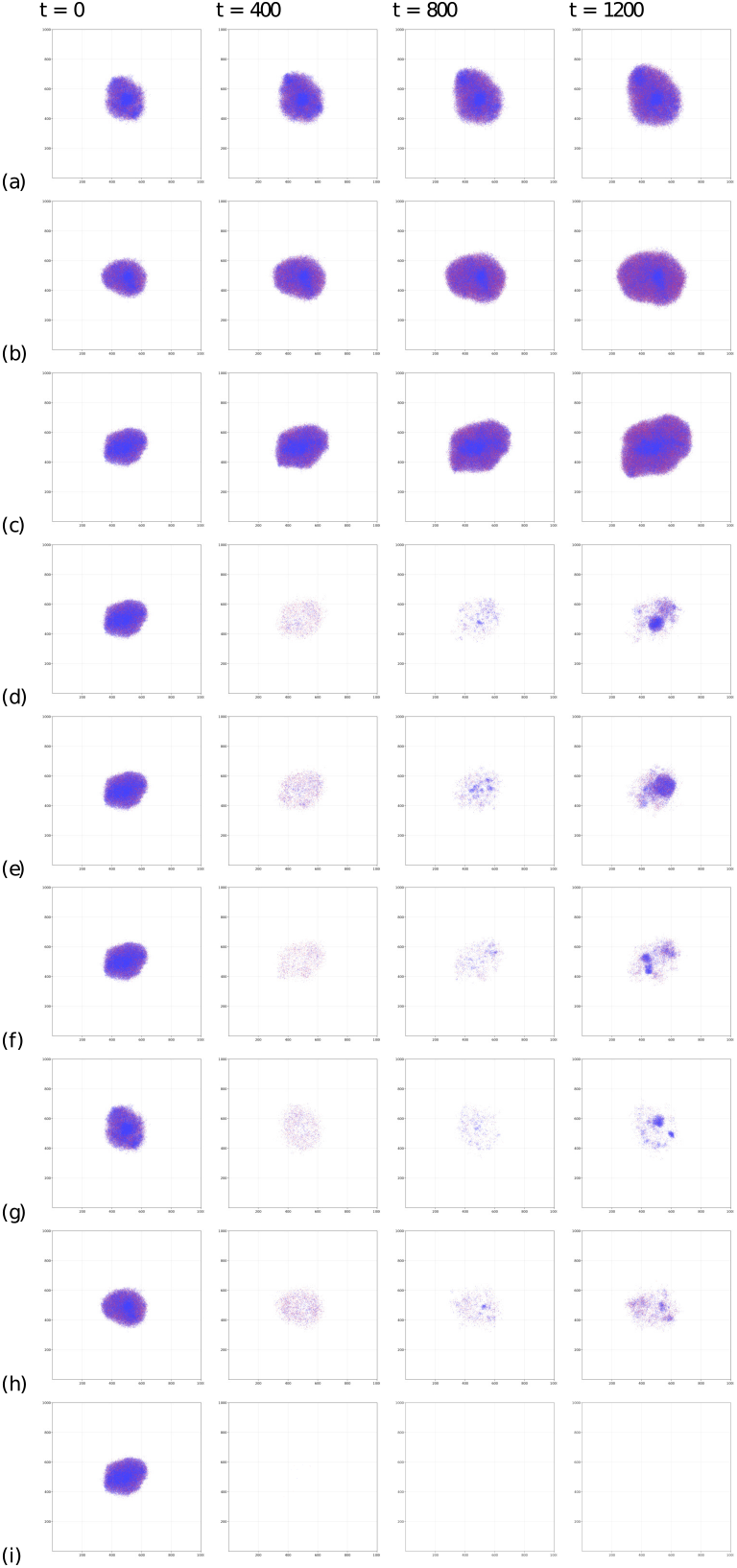
The growth of the tumor at *t* = (0, 400, 800, 1200) from group (a) control *p*_*mut*_ = 0.1, (b) control *p*_*mut*_ = 0.3, (c) control *p*_*mut*_ = 0.5, (d) hyperfractioned radiotherapy *p*_*mut*_ = 0.5, (e) conventional radiotherapy *p*_*mut*_ = 0.5, (f) hypofractioned radiotherapy *p*_*mut*_ = 0.5, (g) conventional radiotherapy *p*_*mut*_ = 0.1, (h) conventional radiotherapy *p*_*mut*_ = 0.3, (i) targeted radiotherapy *p*_*mut*_ = 0.5. Blue and red cells are CCs and CSCs, respectively.

## References

1. Delaney, G., Jacob, S., Featherstone, C. & Barton, M. The role of radiotherapy in cancer treatment. Cancer 104, 1129–1137, DOI: 10.1002/cncr.21324 (2005).

2. Santivasi, W. L. & Xia, F. Ionizing radiation-induced dna damage, response, and repair. Antioxidants & Redox Signal. 21, 251–259, DOI: 10.1089/ars.2013.5668 (2014).

3. Feinendegen, L. E. Reactive oxygen species in cell responses to toxic agents. Hum. & Exp. Toxicol. 21, 85–90, DOI: 10.1191/0960327102ht216oa (2002).

4. Sato, K., Shimokawa, T. & Imai, T. Difference in acquired radioresistance induction between repeated photon and particle irradiation. Front. Oncol. 9, DOI: 10.3389/fonc.2019.01213 (2019).

5. Schulz, A., Meyer, F., Dubrovska, A. & Borgmann, K. Cancer stem cells and radioresistance: DNA repair and beyond. Cancers 11, 862, DOI: 10.3390/cancers11060862 (2019).

6. Arnold, C. R., Mangesius, J., Skvortsova, I.-I. & Ganswindt, U. The role of cancer stem cells in radiation resistance. Front. Oncol. 10, DOI: 10.3389/fonc.2020.00164 (2020).

7. Clarke, M. F. et al. Cancer stem cells—perspectives on current status and future directions: AACR workshop on cancer stem cells. Cancer Res. 66, 9339–9344, DOI: 10.1158/0008-5472.can-06-3126 (2006).

8. Phi, L. T. H. et al. Cancer stem cells (CSCs) in drug resistance and their therapeutic implications in cancer treatment. Stem Cells Int. 2018, 1–16, DOI: 10.1155/2018/5416923 (2018).

9. Charafe-Jauffret, E. et al. Breast cancer cell lines contain functional cancer stem cells with metastatic capacity and a distinct molecular signature. Cancer Res. 69, 1302–1313, DOI: 10.1158/0008-5472.can-08-2741 (2009).

10. Hermann, P. C. et al. Distinct populations of cancer stem cells determine tumor growth and metastatic activity in human pancreatic cancer. Cell Stem Cell 1, 313–323, DOI: 10.1016/j.stem.2007.06.002 (2007).

11. Venkatesan, S., Swanton, C., Taylor, B. S. & Costello, J. F. Treatment-induced mutagenesis and selective pressures sculpt cancer evolution. Cold Spring Harb. Perspectives Medicine 7, a026617, DOI: 10.1101/cshperspect.a026617 (2017).

12. Gray, M. et al. Development and characterisation of acquired radioresistant breast cancer cell lines. Radiat. Oncol. 14, DOI: 10.1186/s13014-019-1268-2 (2019).

13. Bolton, K. L. et al. Cancer therapy shapes the fitness landscape of clonal hematopoiesis. Nat. Genet. 52, 1219–1226, DOI: 10.1038/s41588-020-00710-0 (2020).

14. Marzagalli, M., Fontana, F., Raimondi, M. & Limonta, P. Cancer stem cells—key players in tumor relapse. Cancers 13, 376, DOI: 10.3390/cancers13030376 (2021).

15. Heryanto, Y. D., Achmad, A., Taketomi-Takahashi, A. & Tsushima, Y. In vivo molecular imaging of cancer stem cells. Am. J. Nucl. Med. Mol. Imaging 5, 14–26 (2015).

16. Agliano, A., Calvo, A. & Box, C. The challenge of targeting cancer stem cells to halt metastasis. Semin. Cancer Biol. 44, 25–42, DOI: 10.1016/j.semcancer.2017.03.003 (2017).

17. Enderling, H., Hlatky, L. & Hahnfeldt, P. Migration rules: tumours are conglomerates of self-metastases. Br. J. Cancer 100, 1917–1925, DOI: 10.1038/sj.bjc.6605071 (2009).

18. Poleszczuk, J. & Enderling, H. Cancer stem cell plasticity as tumor growth promoter and catalyst of population collapse. Stem Cells Int. 2016, 1–12, DOI: 10.1155/2016/3923527 (2016).

19. Poleszczuk, J., Hahnfeldt, P. & Enderling, H. Evolution and phenotypic selection of cancer stem cells. PLOS Comput. Biol. 11, e1004025, DOI: 10.1371/journal.pcbi.1004025 (2015).

20. Biava, P. M. et al. Cancer cell reprogramming: Stem cell differentiation stage factors and an agent based model to optimize cancer treatment. Curr. Pharm. Biotechnol. 12, 231–242, DOI: 10.2174/138920111794295783 (2011).

21. Gao, X., McDonald, J. T., Hlatky, L. & Enderling, H. Acute and fractionated irradiation differentially modulate glioma stem cell division kinetics. Cancer Res. 73, 1481–1490, DOI: 10.1158/0008-5472.can-12-3429 (2012).

22. Fukui, R. et al. Tumor radioresistance caused by radiation-induced changes of stem-like cell content and sub-lethal damage repair capability. Sci. Reports 12, DOI: 10.1038/s41598-022-05172-4 (2022).

23. Petersen, C. et al. Repopulation of fadu human squamous cell carcinoma during fractionated radiotherapy correlates with reoxygenation. Int. J. Radiat. Oncol. 51, 483–493, DOI: 10.1016/s0360-3016(01)01686-8 (2001).

24. Allam, A. et al. The effect of the overall treatment time of fractionated irradiation on the tumor control probability of a human soft tissue sarcoma xenograft in nude mice. Int. J. Radiat. Oncol. 32, 105–111, DOI: 10.1016/0360-3016(95)00511-v (1995).

25. Sham, E. & Durand, R. Cell kinetics and repopulation parameters of irradiated xenograft tumours in SCID mice: comparison of two dose-fractionation regimens. Eur. J. Cancer 35, 850–858, DOI: 10.1016/s0959-8049(99)00019-2 (1999).

26. Rofstad, E. K. Repopulation between radiation fractions in human melanoma xenografts. Int. J. Radiat. Oncol. 23, 63–68, DOI: 10.1016/0360-3016(92)90544-r (1992).

27. Fowler, J. F. The phantom of tumor treatment - continually rapid proliferation unmasked. Radiother. Oncol. 22, 156–158, DOI: 10.1016/0167-8140(91)90017-b (1991).

28. Schmidt-Ullrich, R. K. et al. Molecular mechanisms of radiation-induced accelerated repopulation. Radiat. Oncol. Investig. 7, 321–330, DOI: 10.1002/(sici)1520-6823(1999)7:6<321::aid-roi2>3.0.co;2-q (1999).

29. Phillips, T. M., McBride, W. H. & Pajonk, F. The response of cd24 -/low /cd44 + breast cancer-initiating cells to radiation. JNCI: J. Natl. Cancer Inst. 98, 1777–1785, DOI: 10.1093/jnci/djj495 (2006).

30. Bao, S. et al. Glioma stem cells promote radioresistance by preferential activation of the DNA damage response. Nature 444, 756–760, DOI: 10.1038/nature05236 (2006).

31. Gong, L., Zhang, Y., Liu, C., Zhang, M. & Han, S. Application of radiosensitizers in cancer radiotherapy. Int. J. Nanomedicine Volume 16, 1083–1102, DOI: 10.2147/ijn.s290438 (2021).

32. Willers, H., Azzoli, C. G., Santivasi, W. L. & Xia, F. Basic mechanisms of therapeutic resistance to radiation and chemotherapy in lung cancer. The Cancer J. 19, 200–207, DOI: 10.1097/ppo.0b013e318292e4e3 (2013).

33. Stewart, C. A. et al. Single-cell analyses reveal increased intratumoral heterogeneity after the onset of therapy resistance in small-cell lung cancer. Nat. Cancer 1, 423–436, DOI: 10.1038/s43018-019-0020-z (2020).

34. Sprouffske, K. et al. An evolutionary explanation for the presence of cancer nonstem cells in neoplasms. Evol. Appl. 6, 92–101, DOI: 10.1111/eva.12030 (2012).

35. Barker, H. E., Paget, J. T. E., Khan, A. A. & Harrington, K. J. The tumour microenvironment after radiotherapy: mechanisms of resistance and recurrence. Nat. Rev. Cancer 15, 409–425, DOI: 10.1038/nrc3958 (2015).

36. Sherwood, L. M., Parris, E. E. & Folkman, J. Tumor angiogenesis: Therapeutic implications. New Engl. J. Medicine 285, 1182–1186, DOI: 10.1056/nejm197111182852108 (1971).

37. van Diest, P. J. Prognostic value of proliferation in invasive breast cancer: a review. J. Clin. Pathol. 57, 675–681, DOI: 10.1136/jcp.2003.010777 (2004).

38. Martin, B. et al. Ki-67 expression and patients survival in lung cancer: systematic review of the literature with meta-analysis. Br. J. Cancer 91, 2018–2025, DOI: 10.1038/sj.bjc.6602233 (2004).

39. Veronese, S. M. et al. Proliferation index as a prognostic marker in breast cancer. Cancer 71, 3926–3931, DOI: 10.1002/1097-0142(19930615)71:12<3926::aid-cncr2820711221>3.0.co;2-2 (1993).

40. Desai, A., Yan, Y. & Gerson, S. L. Concise reviews: Cancer stem cell targeted therapies: Toward clinical success. Stem Cells Transl. Medicine 8, 75–81, DOI: 10.1002/sctm.18-0123 (2018).

41. Shibata, M. & Hoque, M. O. Targeting cancer stem cells: A strategy for effective eradication of cancer. Cancers 11, 732, DOI: 10.3390/cancers11050732 (2019).

42. Bao, B., Ahmad, A., Azmi, A. S., Ali, S. & Sarkar, F. H. Overview of cancer stem cells (CSCs) and mechanisms of their regulation: Implications for cancer therapy. Curr. Protoc. Pharmacol. 61, DOI: 10.1002/0471141755.ph1425s61 (2013).

43. Hope, K. J., Jin, L. & Dick, J. E. Acute myeloid leukemia originates from a hierarchy of leukemic stem cell classes that differ in self-renewal capacity. Nat. Immunol. 5, 738–743, DOI: 10.1038/ni1080 (2004).

44. Poleszczuk, J., Hahnfeldt, P. & Enderling, H. Biphasic modulation of cancer stem cell-driven solid tumour dynamics in response to reactivated replicative senescence. Cell Prolif. 47, 267–276, DOI: 10.1111/cpr.12101 (2014).

45. Harley, C. B. Telomere loss: mitotic clock or genetic time bomb? Mutat. Res. 256, 271–282, DOI: 10.1016/0921-8734(91)90018–7 (1991).

46. Brú, A. & Casero, D. The effect of pressure on the growth of tumour cell colonies. J. Theor. Biol. 243, 171–180, DOI: 10.1016/j.jtbi.2006.05.020 (2006).

47. Majumdar, S., & Liu, S.-T. Cell division symmetry control and cancer stem cells. AIMS Mol. Sci. 7, 82–101, DOI: 10.3934/molsci.2020006 (2020).

48. Thankamony, A. P., Saxena, K., Murali, R., Jolly, M. K. & Nair, R. Cancer stem cell plasticity – a deadly deal. Front. Mol. Biosci. 7, DOI: 10.3389/fmolb.2020.00079 (2020).

49. Meggiato, T. et al. Spontaneous apoptosis and proliferation in human pancreatic cancer. Pancreas 20, 117–122, DOI: 10.1097/00006676-200003000-00002 (2000).

50. Touil, Y. et al. Colon cancer cells escape 5fu chemotherapy-induced cell death by entering stemness and quiescence associated with the c-yes/YAP axis. Clin. Cancer Res. 20, 837–846, DOI: 10.1158/1078-0432.ccr-13-1854 (2013).

